# A systematic simulation of the effect of salicylic acid on sphingolipid metabolism

**DOI:** 10.1101/011676

**Authors:** Chao Shi, Jian Yin, Zhe Liu, Jian-Xin Wu, Qi Zhao, Jian Ren, Nan Yao

**Affiliations:** State Key Laboratory of Biocontrol, Guangdong Key Laboratory of Plant Resources, School of Life Sciences, Sun Yat-sen University, Guangzhou 510275, P. R. China

**Author notes:** Correspondence: Nan Yao, State Key Laboratory of Biocontrol, Guangdong Key Laboratory of Plant Resources, School of Life Sciences, Sun Yat-sen University, Guangzhou 510275, P. R. China.

**Keywords:** Ceramides_1_, Salicylic acid_2_, Sphingolipid_3_

## Abstract

The phytohormone salicylic acid (SA) affects plant development and defense responses. Recent studies revealed that SA is also involved in the regulation of sphingolipid metabolism, but the details of this regulation remain to be explored. Here, we use *in silico* Flux Balance Analysis (FBA) with published microarray data to construct a whole-cell simulation model, including 23 pathways, 259 reactions and 172 metabolites, to predict the alterations in flux of major sphingolipid species after treatment with exogenous SA. This model predicts significant changes in fluxes of certain sphingolipid species after SA treatment, changes that likely trigger downstream physiological and phenotypic effects. To validate the simulation, we used isotopic non-stationary metabolic flux analysis to measure sphingolipid contents and turnover rate in *Arabidopsis thaliana* seedlings treated with SA or the SA analog benzothiadiazole (BTH). The results show that both SA and BTH affect sphingolipid metabolism by not only concentration of certain species, but also the optimal flux distribution and turnover rate of sphingolipid contents. Our strategy allows us to formally estimate sphingolipid fluxes on a short time scale and gives us a systemic view of the effect of SA on sphingolipid homeostasis.

## INTRODUCTION

Salicylic acid (SA), an important phenolic phytohormone, has well-known roles in pathogen-triggered defense responses including microbe-associated molecular pattern-triggered immunity, effector-triggered immunity, and systemic acquired resistance (Jones and Dangl, 2006; Spoel and Dong, 2012; Yan and Dong, 2014). SA also participates in abiotic stress responses (Vlot *et al.*, 2009; Miura and Tada, 2014) and in plant development, including vegetative and reproductive growth (Vicente and Plasencia, 2011). SA also has indispensible functions in the maintenance of redox homeostasis (Durner and Klessig, 1995 & 1996; Slaymaker *et al.*, 2002) and respiratory pathways (Moore *et al.*, 2002). The SA analog benzothiadiazole (BTH) activates the SA signaling pathway, triggers expression of defense genes (Shimono *et al.,* 2007), and shows similar physiological effects to SA (Lawton *et al.,* 1996).

As a key mediator of defense response, the SA pathway crosstalks with many metabolic pathways. Sphingolipids are a family of complex lipids that have a serine-based head, a fatty acyl chain, and a long-chain base (LCB). Covalent modifications and variability in the length of the fatty acyl chain increase sphingolipid diversity. Sphingolipids are important structural and functional components of the plasma membrane (Hannun and Obeid, 2008) and have important functions in the plant immune response, abiotic stress responses, and developmental regulation (Chen and Cahoon, 2009; Pata *et al.*, 2009; Markham *et al.*, 2013; Bi et al., 2014). In *Arabidopsis*, ceramides, a group of sphingolipids, affect SA-mediated defense responses and programmed cell death (PCD). Some mutants in the sphingolipid metabolic pathway show high levels of expression of defense-related genes, accumulate SA, and undergo PCD. The ceramide kinase deficient mutant *accelerated cell death 5* (*acd5*) accumulates SA and ceramides late in development, but shows increased susceptibility to pathogens (Greenberg *et al.*, 2000; Liang *et al.*, 2003; Bi *et al.*, 2014). Wang *et al.* (2008) reported that the insertion knock out mutant of *Arabidopsis* inositolphosphorylceramide synthase 2 (*erh1*) also spontaneously accumulates SA. Similar increases in SA levels have also been observed in mutants of the sphingosine transfer protein mutant *acd11* (Brodersen *et al.*, 2002), the *Arabidopsis* sphingolipid fatty acid hydroxylase mutants *fah1 fah2* (Konig *et al.*, 2012), and *mips1* (D-myo-inositol 3-phosphate synthase 1) mutants (Meng *et al.*, 2009). Moreover, SA accumulation and PCD signaling mediated by MAPK affect the levels of free LCB (Saucedo-Garcıa *et al.*, 2011). However, *fah1 fah2* mutants accumulate SA and have moderate levels of LCB (Konig *et al.*, 2012). Thus, the SA and sphingolipid pathways have significant but complex crosstalk, particularly in defense and cell death.

Metabolic modeling performs well in prediction of physiological changes and metabolic outcomes resulting from genetic manipulation, where changes in metabolite levels have a strong effect on cellular behavior (Smith and Stitt 2007; Stitt *et al.*, 2010). The genome of *Arabidopsis thaliana* has been sequenced, making whole-genome metabolic reconstruction feasible (Thiele and Palsson, 2010; Seaver *et al.*, 2012). Much of the early modeling work used steady-state Metabolic Flux Analysis (MFA), based on a steady-state model of the plant metabolic network, and on experiments using isotope labeling to trace metabolites of interest (Libourel and Shachar-Hill, 2008; Allen *et al.*, 2009; Kruger *et al.*, 2012). This method provided insights on metabolic organization and modes, but has difficulties in labeling heterotrophic tissues (Sweetlove and Ratcliffe, 2011), over-relies on manual curation of metabolic pathways (Masakapalli *et al.*, 2010; Sweetlove and Ratcliffe, 2011; Kruger *et al.*, 2012), and uses low-throughput detection, making systematic analysis difficult (Lonien and Schwender, 2009; Sweetlove and Ratcliffe, 2011).

By contrast, Flux Balance Analysis (FBA) overcomes many of the drawbacks of MFA. In FBA, a model is established based on a group of ordinary differential equations that formulate a transient quasi-steady state of the metabolic fluxome of target pathways. The duration of the transient flux balance calculated by the FBA model is almost negligible compared to the long-term, fundamental metabolic changes that occur during development or environmental responses (Varma and Palsson, 1994). In addition, FBA does not require isotopic labeling, suits a variety of trophic modes, and is more flexible than steady-state MFA in handling groups of flux distributions by linear programming and other methods for optimization under constraints (Edward and Palsson, 2000; Reed and Palsson, 2003). Several *Arabidopsis* metabolic models based on FBA are available online (Poolman *et al.*, 2009; Dal’Molin *et al.*, 2010; Radrich *et al.*, 2010).

Apart from FBA simulation, fluxomic changes can also be directly measured. Derived from steady-state MFA, isotopic non-stationary metabolic flux analysis (INST MFA) measures *in vivo* time-courses of the transient patterns of isotopic labeling and the steady-state concentrations of various metabolites. Compared with its predecessor, INST MFA has specific advantages, as it is rapid, can deal with monolabeled metabolites (Shastri *et al.*, 2007), can directly measure each flux (Wiechert and Nöh, 2005), and can validate flux predictions made by either laboratory or computational analyses (Nöh and Wiechert, 2006 & 2011; Noack *et al.* 2010).

Since metabolic changes substantially affect the crosstalk between SA and sphingolipids, in this study we constructed a metabolic model to simulate SA-related changes in the sphingolipid pathway. We constructed an *Arabidopsis* whole-cell FBA model including 23 pathways, 259 reactions and 172 metabolites. Based on their relative enrichment and responsiveness to SA stimulation, our model includes 40 sphingolipid species, comprising LCBs, ceramides, hydroxyceramide and glucosylceramides. Due to the lack of flux data on plant sphingolipid metabolism, we used ^15^N labeled INST MFA to measure sphingolipid flux in untreated plants and calibrate the FBA model. After that, additional expression profiles from plants treated with SA and BTH (a SA analog) were supplied to the model. The FBA model was then calculated *in silico* for the prediction and comparison of the optimal flux distribution and flux variability in SA- and BTH-treated and untreated conditions. We then used INST MFA with ^15^N-labeled samples to measure the flux changes directly. Both the computational model and the experiments showed consistent and significant changes in the sphingolipid pathway in response to SA and BTH. Our data gives us a systemic view of the effect of SA on sphingolipid homeostasis.

## MATERIALS AND METHODS

### Plant materials

Wild type *Arabidopsis thaliana* ecotype Columbia seedlings were grown vertically on 1/2x Murashige & Skoog (MS) medium for 10 days after 2-day vernalization. The culture dishes were incubated at 22°C under a 16 h light/8 h dark cycle. For labeling the plant seedlings in liquid medium, the culture dishes were incubated at 22°C with 24 h light.

### Labeling and treatments

The different sphingolipids have many carbon atoms in different positions; therefore, labeling the only nitrogen in the serine-based head group would be much easier for LC-MS/MS measurements. Thus, we used ^15^N serine (Cambridge Isotope Laboratories, Inc. MA, USA) in the labeling experiment. Ten-day-old seedlings were transferred to N-deficient 1/2x MS liquid medium (Yoshimoto *et al.*, 2004) in 12-well culture plates. 5 mM ^15^N-labelled serine was supplied to compensate for the shortage of nitrogen (Hirner *et al.*, 2006) and used as the only source of isotope. For SA and BTH treatments, 100 μM SA or 100 μM BTH was supplied to the labeling medium. The seedlings were treated or not treated for 0, 1, 3, 5, 7, 9, and 24 hours for isotopic non-stationary transient labeling (Nöh and Wiechert, 2006) before sphingolipid extraction.

### Experimental measurement of turnover rate

Since serine has only one nitrogen atom and each sphingolipid has only one serine, the fraction of each labeled sphingolipid species can be simply measured as:

^15^N fraction%=^15^N*100/N

Where ^15^N is the concentration of ^15^N labeled molecules of a specific sphingolipid species, and N is the total concentration of that sphingolipid species, whether labeled or not.

The turnover rate of a sphingolipid species is calculated from the slope of the curve of the time-course of ^15^N incorporation from the initial time that the fraction begins to increase to the fraction stabilizes. Also, the isotopic incorporation rate r can be calculated as:

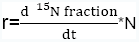

In the measurement, the natural enrichment of ^15^N is relatively constant between samples and treatments.

### Sphingolipid measurements

The plants cultured in labeling medium for the time periods described above were weighed and metabolically quenched by freezing in liquid nitrogen. Sphingolipid species were then extracted and measured by LC-MS/MS as described by Bi *et al.* (2014) with a slight modification to cope with isotopic-labeled sphingolipid species. Major sphingolipid species were subsequently analyzed with a Shimadzu 20A HPLC tandem AB SCIEX TripleTOF 5600^+^ mass spectrometer. The sphingolipid species were analyzed using the software Multiquant (AB SCIEX)

### Metabolic model construction

The *Arabidopsis* whole-cell metabolic model was constructed with 23 pathways, 259 reactions, and 172 metabolites. Primary metabolic pathways refer to the KEGG (Kyoto Encyclopedia of Genes and Genomes http://www.genome.jp/kegg/, Kanehisa et al., 2008), AraCyc database (Mueller *et al.*, 2003), and AraGEM model (Dal’Molin *et al.*, 2010), with manual curation for sphingolipid metabolism, including major ceramide, hydroxyceramide, and glucosylceramide species (Table S1). We used biomass as the objective function with the stoichiometries of major components were assigned to their biomass fraction, which comprises major carbohydrates, amino acids and lipids, according to experiments or the literature (Fiehn *et al.*, 2000; Welti *et al.*, 2002; Dal’Molin *et al.*, 2010). For sphingolipid species, the objective function stoichiometries were set to the adjusted isotopic incorporation rate in labeling experiments.

### Flux balance analysis (FBA)

Flux balance modeling uses a group of ordinary differential equations. The analysis requires a stoichiometric matrix (S) and a vector (v) built for each reaction, where s_ij_ in the S matrix is the stoichiometric number of the ith metabolite in the jth reaction and v_j_ is the rate of the jth reaction, which is subjected to upper and lower boundary constraints. To reach the *in silico* “quasi-steady state”, the following condition must be fulfilled:

S⋅*v*=0

After solving the FBA equation with the constraints above (Edwards and Palsson, 2000; Edwards *et al.*, 2001), a linear-programming optimization method (Edwards and Palsson, 2000) was applied to pick the most plausible (groups of) flux distributions among the solution space according to the objective setting.

### *in silico* SA and BTH treatments

To incorporate the effect of exogenous SA and BTH on the wild-type plant into the model, we used published microarray data for SA- and BTH-treated *Arabidopsis* (for SA, Leeuwen *et al.*, 2007; for BTH, Wang *et al.*, 2006). We assumed that the metabolic flux changed following the same trend as the respective gene expression levels. Therefore, when genes were matched to microarray probes to identify their changes in expression following each treatment, we picked genes that changed more than 1.5-fold in SA-treated plants and more than 2-fold in BTH-treated plants (Table S2). Then, the adjusted model was recalculated for optimal flux distribution.

### Flux variability analysis (FVA)

The stoichiometry model is a self-balancing model in that any flux distributions that fulfill the constraints are involved in its solution space. Through the sampling of the solution space or sensitivity analysis, each reaction is tested for its possible upper flux limit and lower flux limit under constraints (Mahadevan and Schilling, 2003). The calculated range of each flux is an important indicator of the role of the corresponding reaction in the robustness of the whole network. To make a physiologically relevant estimation, we sample the flux space which achieves at least 80% of optimal objective rate (in our model, the biomass production) under untreated or treated condition.

### Simulation environment

The model of *Arabidopsis* was built in SBML (Systems Biology Makeup Language) (Hucka *et al.*, 2003) in XML format. SBML Toolbox 2.0.2 (Keating *et al.*, 2006; Schmidt and Jirstrand, 2006) and COBRA Toolbox 2.0.5 (Schellenberger *et al.*, 2011) in MATLAB 2012a (Mathworks Inc.; Natick, MA) were used for model construction and calculation. Linear programming was performed with GLPK (GNU Linear Programming Kit, http://www.gnu.org/software/glpk/). The rank-test and multiple covariance analysis are performed using IBM SPSS Statistics 19 (IBM Corp. Released 2010. IBM SPSS Statistics for Windows, Version 19.0. Armonk, NY: IBM Corp.).

## RESULTS

### Model construction for plant sphingolipid metabolism

We aimed to explore the changes in plant sphingolipid metabolism in response to SA, by using computational modeling and experiments. Although sphingolipids function as important components in plant development and stress responses, their metabolism remains obscure, with few measured network parameters. FBA is well suited to the simulation of a metabolic fluxome with poorly-understood dynamics (Varma and Palsson, 1994), as FBA requires only the stoichiometric relationship in each reaction and the objective function for optimization. In our model, the numbers of molecules of reactants and products in known reactions were obtained from public databases (see Materials and Methods). For the sphingolipid pathways (Table S1), those reactions that have not been determined were inferred from their atomic composition or similar reactions. Considering that metabolic balances are mainly affected by a few metabolites that are either in a hub of the network or have high turnover, we picked the sphingolipid species that are relatively abundant or central to the known network (Table 1). Since inositolphosphorylceramide and its derivatives are difficult to measure in plants, we excluded those species from our model.

### Isotopic non-stationary transient labeling of sphingolipids

To inform the objective function and to validate the model’s prediction, we used the *in vivo* fluxomic method of isotopic non-stationary transient labeling (INST) to directly measure the turnover rate of plant sphingolipids. In previous work, ^13^C INST was mostly used to examine central pathways such as glucose metabolism or photosynthesis (Noack *et al.*, 2010; Nöh and Wiechert, 2011), where limited numbers of labeled fragments are detected by mass spectrometry. However, the simplest sphingolipid has at least 18 carbon atoms, and their combined transitions, modifications and fragmentation would generate large numbers of labeled fragments; therefore mass spectrometry quantification of ^13^C labeled sphingolipid would be extremely difficult, whatever the labeling material used. Therefore, we selected the only nitrogen atom in the head of each sphingolipid as the labeling position. To distinguish between artificial and natural ^15^N, we measured the composition of natural ^15^N sphingolipid in unlabeled samples, finding different levels of natural ^15^N in each sphingolipid species. This fraction is constant between measurements and treatments in each species, and thus cannot affect the comparison of isotopic incorporation rates between experiments.

We transiently labeled 10-day-old seedlings in a time course. The isotopic incorporation curves (see representative species shown in Figure 1) reveal that the labeled serine is absorbed and incorporated into sphingolipid in the first hour of labeling, followed by a uniform turnover rate. For LCB (Figure 1D), ceramide (Figure 1A), and hydroxyceramide species (Figure 1B), the isotopic incorporation curves gradually flatten and finally reach an isotopic fraction balance between 9 and 24 h. A noticeable, small drop occurs around the 5^th^ hour of incorporation in LCB (Figure 1D). The incorporation of ^15^N in these simple sphingolipids is fast, and the final balanced isotopic fraction can reach 40-65% (Figure 1A, 1B and 1D). By contrast, between 9 and 24 h, the labeled fraction constantly rose for the glucosylceramides (Figure 1C), which had a lower rate of incorporation than the ceramides or hydroxyceramides. Combined with the concentration of sphingolipids, we calculated the isotopic incorporation rate as shown in Table 2.

**Figure 1.**
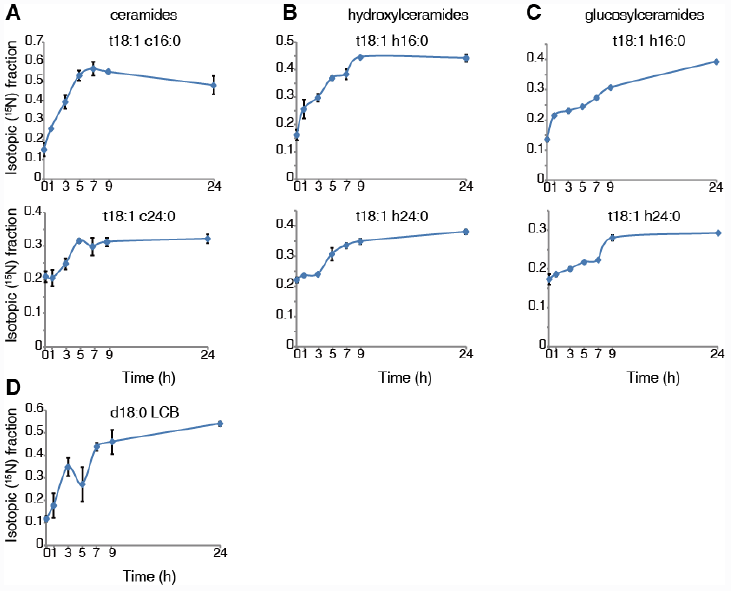
^15^N incorporation curves for sphingolipid species. Ten-day-old wild-type seedlings were transferred to 5 mM ^15^N- serine labeled N-deficient 1/2x MS liquid medium for the indicated times. Sphingolipids were then extracted and measured as described in Methods. The ^15^N fraction incorporation curve was calculated based on the formula shown in Methods. Error bars represent the means ±SE from triplicate biological repeats. The measured sphingolipid species were: ceramide (A), hydroxyceramide (B), gluocosylceramide (C) and LCB (D). LCB and fatty acid in ceramide species represent: LCB; d/t (di/trihydroxy) 18 (18 carbon chain): 1 (one desaturation) followed by fatty acid; c/h/g (non-hydroxyl / hydroxyl / glucosy and hydroxyl) 24 (24 carbon chain): 0 (no desaturations).

### Flux balance analysis (FBA) of the flux distribution in untreated plants

The objective function in the FBA model guides flux determination by simulating a transient flux distribution. However, biomass at a certain time is the complex result of development through the organism’s entire life, and hence cannot be a relevant principle in setting the objective function in our model of the *Arabidopsis* seedling. Instead, the objective function stoichiometries of the sphingolipid pathway was built and adjusted from isotopic incorporation rates in the above labeling experiments (Table 1). Then, flux balance optimization was performed. Figure 2 shows the simulated flux distributions of sphingolipid species in untreated plants.

**Figure 2.**
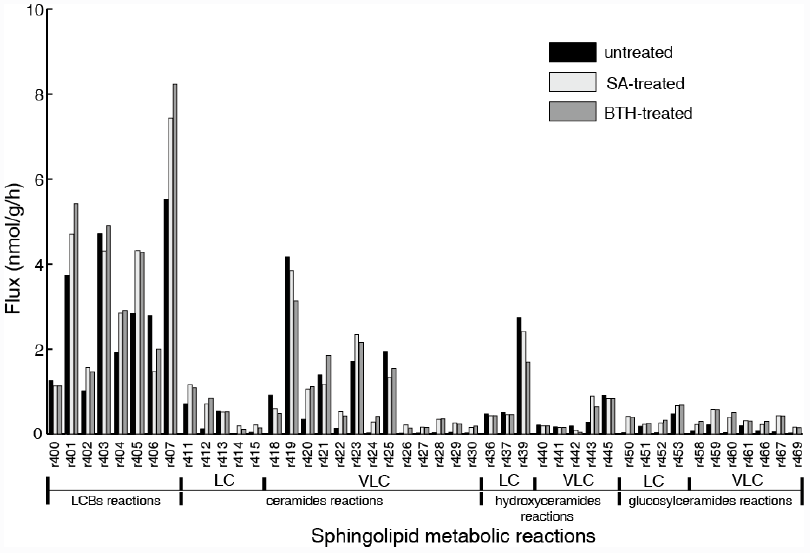
Simulated flux distribution of selected sphingolipid species. The untreated plants (black) and *in silico* SA (light gray) and BTH-treated plants (gray) were taken from the flux balance model. The effects of exogenous SA and BTH were simulated by changing the target flux bound proportional to its related gene expression alteration identified by published microarray data (Wang *et al.*, 2006; van Leeuwen *et al.*, 2007). LC, long-chain (≤C18); VLC: very-long-chain (>C18).

**Table 1.** Overview of sphingolipid species in the FBA model.

| Symbol | Sphingolipid species | Pool size<br>(nmol·g<sup>-1</sup>) | Stoichiometry in<br>objective function |
| --- | --- | --- | --- |
| d18:0 LCB | Long-chain base | 1.686182 | 0.050201 |
| d18:1 LCB | Long-chain base | 0.323814 | 0.017119 |
| t18:0 LCB | Long-chain base | 2.243848 | 0.044619 |
| t18:1 LCB | Long-chain base | 0.894187 | 8.05E-05 |
| t18:1 c16:0 | long-chain ceramide | 1.375136 | 0.14095 |
| t18:0 c16:0 | long-chain ceramide | 0.078273 | 0.006289 |
| d18:1 c16:0 | long-chain ceramide | 0.103578 | 0.017411 |
| d18:0 c16:0 | long-chain ceramide | 0.323513 | 0.040446 |
| t18:0 c24:0 | very-long-chain ceramide | 17.51997 | 0.47712 |
| t18:1 c24:0 | very-long-chain ceramide | 30.1346 | 0.775466 |
| t18:0 c24:1 | very-long-chain ceramide | 4.702168 | 0.119545 |
| t18:1 c24:1 | very-long-chain ceramide | 10.12495 | 0.344293 |
| t18:0 c26:0 | very-long-chain ceramide | 5.964148 | 0.129493 |
| t18:1 c26:0 | very-long-chain ceramide | 29.47465 | 0.671015 |
| t18:0 c26:1 | very-long-chain ceramide | 0.326354 | 0.005744 |
| t18:1 c26:1 | very-long-chain ceramide | 6.565916 | 0.208064 |
| t18:1 h160 | long-chain hydroxyceramide | 6.406314 | 0.154383 |
| t18:0 h160 | long-chain hydroxyceramide | 0.682043 | 0.012748 |
| d18:1 h16:0 | long-chain hydroxyceramide | 0.351323 | 0.020931 |
| d18:0 h16:0 | long-chain hydroxyceramide | 0.292355 | 0.019623 |
| t18:0 h24:0 | very-long-chain hydroxyceramide | 10.38919 | 0.01712 |
| t18:1 h24:0 | very-long-chain hydroxyceramide | 80.91966 | 1.148618 |
| t18:0 h24:1 | very-long-chain hydroxyceramide | 8.615409 | 0.124845 |
| t18:1 h24:1 | very-long-chain hydroxyceramide | 0.169527 | 1.53E-05 |
| t18:0 h26:0 | very-long-chain hydroxyceramide | 3.30798 | 0.003149 |
| t18:1 h26:0 | very-long-chain hydroxyceramide | 17.71101 | 0.218833 |
| t18:0 h26:1 | very-long-chain hydroxyceramide | 1.005991 | 9.05E-05 |
| t18:1 h26:1 | very-long-chain hydroxyceramide | 10.14596 | 0.27478 |
| t18:1 h16:0 | long-chain glucosylceramide | 7.336978 | 0.03589 |
| t18:0 h16:0 | long-chain glucosylceramide | 0.00001 | 9.00E-10 |
| d18:1 h16:0 | long-chain glucosylceramide | 23.26684 | 0.177984 |
| d18:0 h16:0 | long-chain glucosylceramide | 0.191598 | 0.001506 |
| t18:0 h24:0 | very-long-chain glucosylceramide | 1.55239 | 0.00014 |
| t18:1 h24:0 | very-long-chain glucosylceramide | 14.59155 | 0.055296 |
| t18:0 h24:1 | very-long-chain glucosylceramide | 0.00001 | 9.00E-10 |
| t18:1 h24:1 | very-long-chain glucosylceramide | 17.28822 | 0.057862 |
| t18:0 h26:0 | very-long-chain glucosylceramide | 0.00001 | 9.00E-10 |
| t18:1 h26:0 | very-long-chain glucosylceramide | 8.470761 | 0.032563 |
| t18:0 h26:1 | very-long-chain glucosylceramide | 0.00001 | 9.00E-10 |
| t18:1 h26:1 | very-long-chain glucosylceramide | 5.706558 | 0.016164 |

The simulation data in Figure 2 shows that LCBs, very-long-chain ceramides and hydroxyceramides compose the highest fraction of total flux. The ^15^N labeled INST MFA is consistent with these simulated results for the measured fast isotopic incorporation and high fraction of stabilized isotopic final level of LCB, ceramides and hydroxyceramides (Figure 1). These results demonstrate that LCBs, the sphingolipids that have the smallest pool size, also have the highest turnover among plant sphingolipids. Very-long-chain ceramides and hydroxyceramides are important not only for their hub position connecting glucosylceramides and sphingosine, but also carry a huge flux throughput in sphingolipid turnover and thus help maintain sphingolipid homeostasis. Both the simulation and experimental results indicate that these sphingolipid species are probably more responsive to disturbance, and thus are frequently used by pathogens to manipulate or interfere with host sphingolipid metabolism (Asai *et al.*, 2000; Markham *et al.*, 2011, Bi *et al*., 2014).

Although the glucosylceramides have much larger pool sizes (Table 1) than the ceramides, hydroxyceramides, or LCBs, they have smaller metabolic fluxes than their precursors (Figure 2). These results are validated by the slow but lasting incorporation of isotope into glucosylceramide pools (Figure 1C). The relatively slow turnover is in accordance with the function of glucosylceramides as membrane structural components, indicating a slow but continuous accumulation in the cell membrane during plant development. The accordance of simulation and experiment results also supports our choice of objective function stoichiometries setting, for the scale of simulated and measured sphingolipid metabolic flux distribution (Figure 2 and Table 2) is nearly unrelated to the distribution of sphingolipid biomass (Table 1).

**Table 2.**
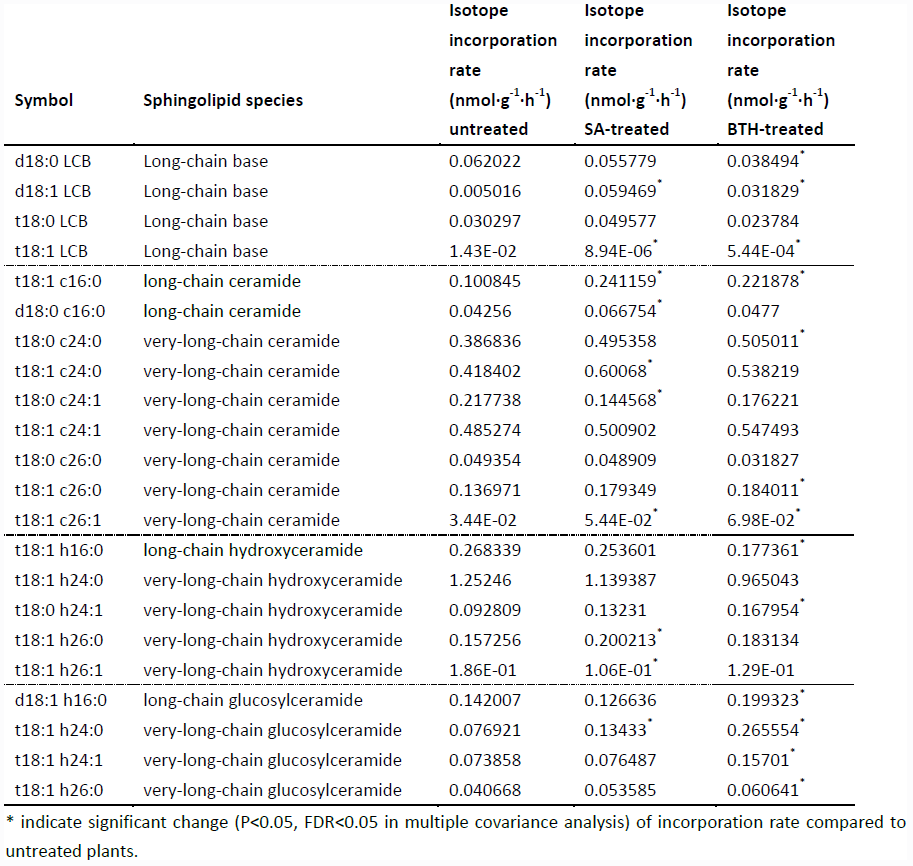
Isotopic incorporation rate for major sphingolipids, with or without 100 M SA or 100 M BTH treatments

| Symbol | Sphingolipid species | Isotope<br>incorporation<br>rate<br>($\text{nmol}\cdot\text{g}^{-1}\cdot\text{h}^{-1}$) | Isotope<br>incorporation<br>rate<br>($\text{nmol}\cdot\text{g}^{-1}\cdot\text{h}^{-1}$) | Isotope<br>incorporation<br>rate<br>($\text{nmol}\cdot\text{g}^{-1}\cdot\text{h}^{-1}$) |
| --- | --- | --- | --- | --- |
|  |  | untreated | SA-treated | BTH-treated |
| d18:0 LCB | Long-chain base | 0.062022 | 0.055779 | 0.038494* |
| d18:1 LCB | Long-chain base | 0.005016 | 0.059469* | 0.031829* |
| t18:0 LCB | Long-chain base | 0.030297 | 0.049577 | 0.023784 |
| t18:1 LCB | Long-chain base | 1.43E-02 | 8.94E-06* | 5.44E-04* |
| t18:1 c16:0 | long-chain ceramide | 0.100845 | 0.241159* | 0.221878* |
| d18:0 c16:0 | long-chain ceramide | 0.04256 | 0.066754* | 0.0477 |
| t18:0 c24:0 | very-long-chain ceramide | 0.386836 | 0.495358 | 0.505011* |
| t18:1 c24:0 | very-long-chain ceramide | 0.418402 | 0.60068* | 0.538219 |
| t18:0 c24:1 | very-long-chain ceramide | 0.217738 | 0.144568* | 0.176221 |
| t18:1 c24:1 | very-long-chain ceramide | 0.485274 | 0.500902 | 0.547493 |
| t18:0 c26:0 | very-long-chain ceramide | 0.049354 | 0.048909 | 0.031827 |
| t18:1 c26:0 | very-long-chain ceramide | 0.136971 | 0.179349 | 0.184011* |
| t18:1 c26:1 | very-long-chain ceramide | 3.44E-02 | 5.44E-02* | 6.98E-02* |
| t18:1 h16:0 | long-chain hydroxyceramide | 0.268339 | 0.253601 | 0.177361* |
| t18:1 h24:0 | very-long-chain hydroxyceramide | 1.25246 | 1.139387 | 0.965043 |
| t18:0 h24:1 | very-long-chain hydroxyceramide | 0.092809 | 0.13231 | 0.167954* |
| t18:1 h26:0 | very-long-chain hydroxyceramide | 0.157256 | 0.200213* | 0.183134 |
| t18:1 h26:1 | very-long-chain hydroxyceramide | 1.86E-01 | 1.06E-01* | 1.29E-01 |
| d18:1 h16:0 | long-chain glucosylceramide | 0.142007 | 0.126636 | 0.199323* |
| t18:1 h24:0 | very-long-chain glucosylceramide | 0.076921 | 0.13433* | 0.265554* |
| t18:1 h24:1 | very-long-chain glucosylceramide | 0.073858 | 0.076487 | 0.15701* |
| t18:1 h26:0 | very-long-chain glucosylceramide | 0.040668 | 0.053585 | 0.060641* |

\* indicate significant change ( $P < 0.05$ , $\text{FDR} < 0.05$ in multiple covariance analysis) of incorporation rate compared to untreated plants.

### *In silico* SA and BTH treatments

The FBA model hypothesizes the quasi-steady state condition of the target network, and we assume that the sphingolipid pathway will reach at least a transient metabolic balance after SA treatment. Thus, we employed the previous model simulating the resting state to predict the effects of SA treatment. We first used data from microarray analysis of SA- and BTH-treated plants to simulate the effect of these treatments on sphingolipid flux. Sphingolipid-related genes were chosen (see Method) from two microarrays (Table S2). *LAG 1 HOMOLOG 2* (*LOH2*), which encodes a ceramide synthase (Brandwagt *et al.,* 2000; Ternes et al., 2011), shows the highest up-regulation after both SA and BTH treatments, and other genes show different expression between treatments (*SPHINGOID BASE HYDROXYLASE 2* (*SBH2*), *FATTY ACID/SPHINGOLIPID DESATURASE* (*SLD*), *FATTY ACID HYDROXYLASE 2* (*FAH2*), *SPHINGOSINE-1-PHOSPHATE LYASE* (*AtDPL1*). The reactions regulated by the genes with altered transcript levels were then picked for incorporation in the model. The flux boundaries of these reactions were altered based on the gene expression level, and the adjusted model was recalculated for flux balance analysis.

Compared with the model simulating the resting state, *in silico* SA and BTH treatments resulted in a nearly three-fold increase of flux in long-chain ceramide species (Figure 2), which is consistent with the up-regulation of *LOH2* in the microarray data. In particular, simulated SA and BTH treatment both showed a significant rise in metabolism of trihydroxy glucosylceramides. This increase is not specific to fatty acid species, which showed an increase in both trihydroxy long-chain and very-long-chain glucosylceramides (Figure 2). These results are consistent with the data from ^15^N labeled INST MFA (Table 2). Interestingly, the microarray data showed no significant changes in genes that directly catalyze the pathways in glucosylceramide metabolism, nor any related to glucosylceramide, in response to SA or BTH treatment (Table S2). Considering the down-regulation of *SBH2* under BTH treatment, we believe that the increase of glucosylceramide metabolism may mainly be induced by the upstream up-regulation of *LOH2*. Since the increase of the turnover rate is not linked to metabolite concentration, the changes of glucosylceramides are almost negligible by typical quantitative LC-MS/MS measurement, but the increase in lipid renewal may have indispensible functions in the sensitivity of membrane-based cell signaling.

In this simulation, although some genes change differently in response to SA and BTH treatment (Table S2), they have similar effects on sphingolipid metabolism. Our model also proposes a possible mechanism by which BTH affects the network under flux balance constraint without mimicking all the gene expression changes of its counterpart.

### Isotopic non-stationary transient measurement of the effect of SA and BTH

Last, to confirm the predictions of the model, we directly measured the *in vivo* flux change in response to SA and BTH treatments. For SA and BTH treatments, the isotope incorporation rate significantly increased for certain sphingolipid species such as LCBs and ceramides (Table 2). These results are consistent with our FBA model (Figure 2).

### Flux variability analysis

To examine the change in network rigidity in response to SA and BTH treatments, we estimated the accessible flux ranges of sphingolipid species *in silico*. To make a physiologically relevant estimation, we sampled the flux space which achieves at least 80% of optimal objective rate (in our model, the biomass production) under untreated or treated condition. We sorted the flux range into three types (Oberhardt *et al.*, 2010): rigid flux (flux range near zero but with non-zero flux value), bounded flexible flux, and infinitely flexible flux (boundary spans from 0 or -1000 to 1000 in the model). In the fluxome of treated and untreated plants, LCB fluxes were infinitely flexible, showing a high capacity to tolerate disturbance, ceramide and glycosylceramide fluxes showed bounded flexibility, and hydroxyceramide fluxes were rigid (Table S3). The limited flux variability of most sphingolipids is consistent with stoichiometric modeling result in *S. cerevisiae* (Ozbayraktar and Ulgen, 2011). Similar with isotopic incorporation experiments, we found the disturbances of flux variability in the reaction of ceramide and glucosylceramide metabolic fluxes after SA and BTH treatments (Figure 3), indicating freedom of adjusting their metabolism under the prerequisite of sphingolipid flux homeostasis during defense process.

**Figure 3.**
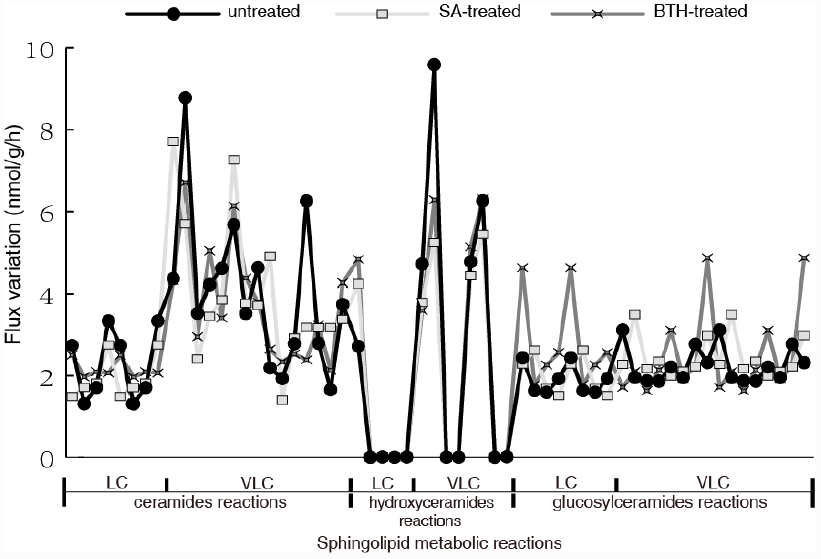
Flux variations among selected reactions of different sphingolipid species under no treatment or *in silico* SA or BTH treatments. LCB fluxes are omitted for their too large flux variation up to 1000 nmol/g/h, while other fluxes keep rigid after *in silico* treatment (see supplemental Table S3).

## DISCUSSION

Our FBA model and isotope labeling experiments systematically explored the alterations in the sphingolipid pathway that occur in response to SA and BTH. It is well-known that traditional metabolic responses are often considered to be significant changes of certain metabolites concentration. However, the systematic responses caused by plant activator and phytohormone cannot be achieved by only doubling the concentration fold of certain nodes without affecting the dynamic properties of the whole network. To panoramically detect these underlying changes of network parameters presented by up and down of certain nodes, one of the most direct measurements is the fluxome. FBA analysis has been applied in microbial metabolic engineering and modeling of other systems. However, construction of the model for sphingolipid metabolism presented difficulties related to the unique features of sphingolipid pathways. Although sphingolipid species are among the most reactive components in plant development and stress responses, they reside in the periphery of the network of plant metabolism, having loose metabolic connections with other subnetworks. Their lack of connection and remote position make the flux in the self-balanced function more susceptible to the objective settings, rather than being affected by artificial constraints and neighboring subnetworks.

Indeed, there are other studies concerning sphingolipids in *S. cerevisiae* (Ozbayraktar and Ulgen, 2011) where the sphingolipid pathways are also remote from central metabolism, but these models are backed by experimental data on enzyme kinetic parameters or known fluxes. In experiments, plant sphingolipid pathways are difficult to explore because of their vast diversity, low abundance, and lack of sensitive and replicable measurements. In addition, the enzymes linking metabolites often are embedded in the layers of membranes, making the isolation and estimation of their kinetic properties difficult. Until now, a limited set of experiments has determined only a rough scheme of plant sphingolipid metabolism. Considering that, we used the experimentally measured isotopic incorporation rate to set the stoichiometry of each sphingolipid species as stoichiomeries in the objective function, and we found that the resulting flux distribution of each species is in accordance with isotopic incorporation pattern, demonstrating that isotopic incorporation data produces a better fit than biomass fraction in objective stoichiomery determination, as the maximization of biomass is often considered as the aim of plant metabolism regardless of any inconsistency between biomass contents and generation rate of each component.

In the experimental part, isotopic transient labeling provided a direct measurement of *in vivo* flux. We note that none of the sphingolipid species reached 100% labeled. Similar phenomena were also observed in other experiments (Delwiche and Sharkey, 1993; Hasunuma *et al.*, 2010). Considering the internal serine sources and anaplerotic reactions of complex existing sphingolipids, the pattern indicates a balance of labeled and unlabeled sphingolipids in the metabolic pool. Since the only exogenous source of nitrogen is labeled, we can also speculate on the utilization of external and internal sources of nitrogen in sphingolipid synthesis from the isotopic incorporation curve.

There are various models linking plant sphingolipid pathways with hormones and their synergistic role in plant development and stress responses. In these models, the possible sphingolipid inducers of defense responses include LCBs (Saucedo-Garcıa *et al.*, 2011) and ceramides (Markham *et al.*, 2011, Bi *et al.,* 2014), with SA both up- and downstream of sphingolipid-mediated PCD (Saucedo-Garcıa *et al.*, 2011; Bi *et al*., 2014). As sphingolipid mutants often accumulate SA, the effect of SA on ceramide species may include positive feedback on the imbalance of sphingolipids. Our results are in accordance with the observed frequent variation in the concentration of LCB and sometimes ceramide, but less variation in the concentrations of hydroxyceramide and glucosylceramide in wild-type *Arabidopsis*. Functionally speaking, since LCB and ceramides are fundamental to sphingolipid metabolism and show high flux flexibility, they can be more responsive to stimuli such as SA or BTH without disrupting the total fluxomic balance of sphingolipid metabolism

In a living cell, the synthesis and degradation of all substances occurs through metabolism. However, current research tends to separate metabolites and functional molecules. The most exciting aspect of plant sphingolipids is that they are themselves metabolites and functional molecules. Our current model only deals with their metabolic properties in a self-balanced manner. It will be interesting to incorporate the signaling network that involves sphingolipids to build an integrated model to consider the direct effect of metabolism on cell signaling.

## CONCLUSION

In this study, we established a sphingolipid FBA model and used ^15^N labeled isotopic transient labeling to systematically explore the effects of SA and BTH on sphingolipid metabolic pathways. The results show increases in ceramide and glucosylceramide flux in response to exogenous SA and BTH and also alteration of their flux variability. Our results also give us insights that help explain the mechanism of crosstalk between SA and sphingolipids, and their roles in the plant defense response.

## ACKNOWLEDGEMENT

We thank members in the Yao and Ren laboratories for assistance with this work. This work was supported by the National Key Basic Science 973 Program (2012CB114006), National Natural Science Foundation of China (31170247), and the Fundamental Research Funds for the Central Universities (13lgjc27).

## Supplemental information

**Table S1.** The indexes, categories and equations of sphingolipid–related reactions in our FBA model.

**Table S2.** Sphingolipid-related gene expression changes in SA- and BTH-treated plants from microarrays data published by van Leeuwen *et al.* (2007) and Wang *et al*. (2006).

**Table S3.** Simulated flux variability of sphingolipid-related reactions in untreated and *in silico* SA- or BTH- treated plants.

